# Integrated evaluation of immune system perturbation using structural, functional and cellular immunotoxicity endpoints in rats

**DOI:** 10.64898/2026.03.22.713556

**Authors:** Vinay Lomash, M Srinivasan, P Mahesh, Sayeed Azgar, Gayathri Venkatesan, Babitha Joseph

## Abstract

Evaluation of unintended immunotoxicity represents an important component of nonclinical safety assessment, as perturbation of immune function may increase susceptibility to infection, impair vaccine responses, and disrupt immune homeostasis. Regulatory guidance, including the ICH S8 Immunotoxicity Guideline, recommends a weight-of-evidence approach in which observations from conventional toxicological endpoints are integrated with functional immune assays to support interpretation of immune system effects. The present study applied an integrated immunotoxicity evaluation framework to examine concordance among structural, functional, and cellular immune endpoints in male Sprague–Dawley rats using a well-characterized immunosuppressive reference compound. Hematological evaluation revealed leukopenia characterized primarily by lymphocyte depletion. Reductions in spleen and thymus weights were accompanied by histopathological evidence of lymphoid depletion in multiple immune tissues, including spleen, thymus, lymph nodes, Peyer’s patches, and bone marrow. Functional immune competence was assessed through hemagglutination antibody response to sheep red blood cells and delayed-type hypersensitivity assays, both of which demonstrated marked suppression of adaptive immune responses. Flow cytometric immunophenotyping further demonstrated substantial reductions in B-cell populations and decreases in CD4⁺ and CD8⁺ T-cell counts, whereas NK cell populations were comparatively less affected. The concordance of hematological alterations, lymphoid tissue changes, impaired functional immune responses, and lymphocyte subset depletion provides integrated evidence of immune system perturbation. These findings demonstrate that complementary immunotoxicity endpoints collectively support hazard characterization of immune system effects under GLP conditions.

**Graphical Abstract:** 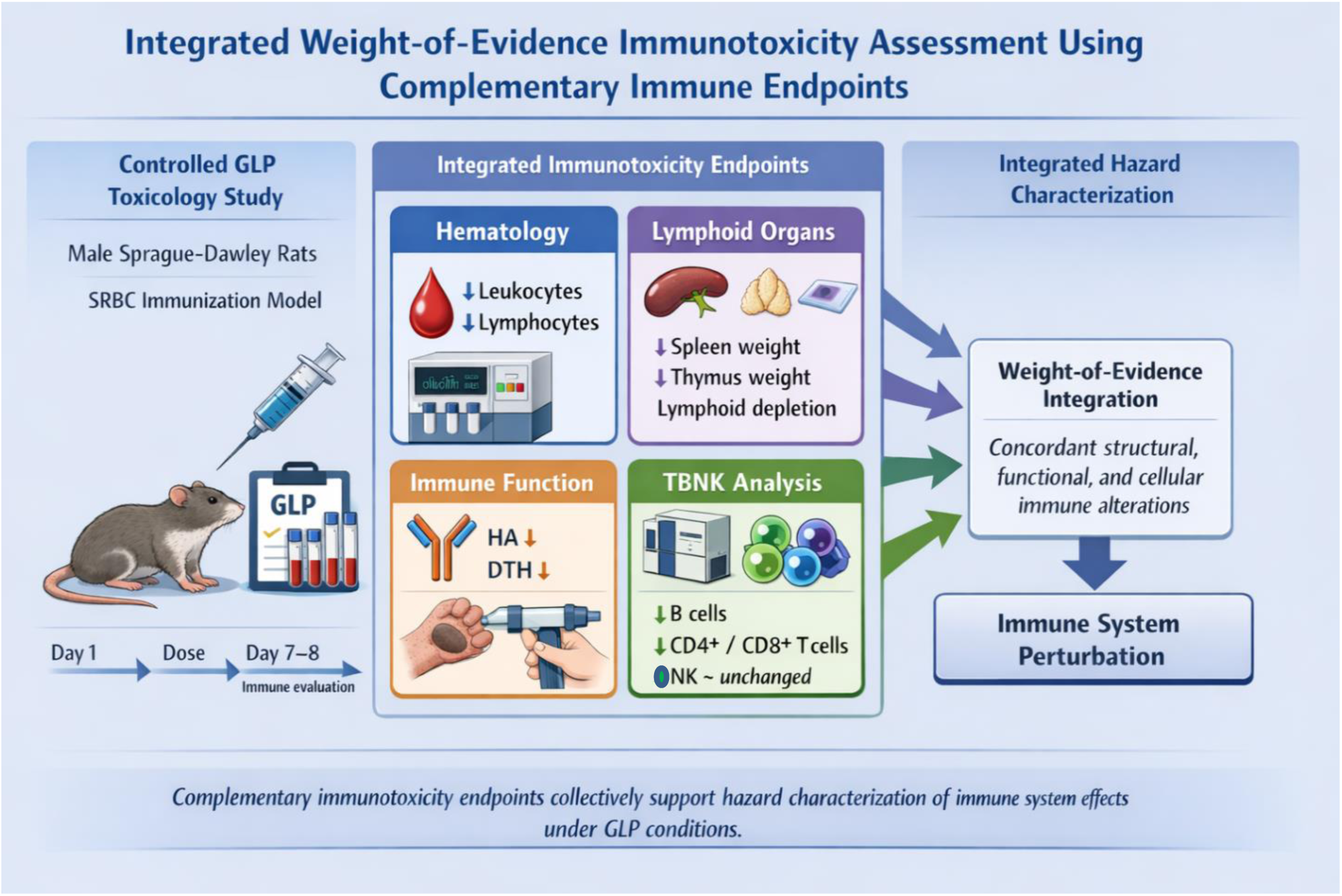

## Introduction

The immune system plays a central role in host defense, maintenance of tissue homeostasis, and surveillance against infectious and neoplastic diseases. Exposure to pharmaceuticals or other xenobiotics capable of altering immune function may lead to increased susceptibility to infections, impaired vaccine responses, dysregulated inflammatory processes, or altered immune surveillance. Consequently, evaluation of unintended immunotoxicity has become an important component of nonclinical safety assessment during development of pharmaceuticals as well as certain industrial and environmental chemicals.

Regulatory authorities have established guidance frameworks to support systematic identification of compounds that may adversely affect immune competence. The ICH S8 Guideline recommends a risk-based strategy in which potential immunotoxic effects are first identified through findings in standard repeat-dose toxicity studies, including alterations in hematology, lymphoid organ weights, and histopathology^1^. Where such observations or pharmacological considerations indicate possible immune involvement, targeted investigations may be undertaken to evaluate functional immune responses. Similarly, the EPA OPPTS 870.7800 Guideline^2^ outlines approaches for assessing immune competence in regulatory toxicology programs intended to detect substances capable of altering immune responses. Within these regulatory frameworks, interpretation of immunotoxicity data generally follows a weight-of-evidence approach in which multiple lines of evidence are considered collectively during hazard identification and risk characterization^3,4^.

The scientific basis for these regulatory approaches derives from early experimental investigations that evaluated the predictive value of specific immune function assays. Studies conducted by Luster and colleagues demonstrated that a limited set of immune endpoints including the T-dependent antibody response (TDAR), natural killer (NK) cell activity, and delayed-type hypersensitivity (DTH) exhibit high sensitivity and concordance in detecting compounds with immunotoxic potential^5,6^. These investigations contributed to the development of tiered testing strategies in which initial signals observed in conventional toxicity parameters can be further evaluated using functional assays that measure adaptive or innate immune competence. Subsequent work has continued to emphasize the importance of integrating functional immune assays with traditional toxicological endpoints in order to improve interpretation of immune-related findings within safety assessment^7^.

From a regulatory toxicology perspective, evaluation of immunotoxicity generally progresses through sequential interpretive stages. Initial hazard identification is based on alterations in leukocyte populations, changes in lymphoid organ weights, and histopathological findings in immune tissues. Where evidence of immune modulation is observed, functional confirmation may be obtained through assays that assess adaptive immune responses, including TDAR or cellular immune responses such as DTH. In some cases, mechanistic characterization may also be undertaken through immunophenotyping or cellular subset analysis to support biological plausibility and strengthen interpretation of the observed effects^1,4^.

Although guidance documents detail multi-test batteries and correlations of immunotoxicity assays, most immunotoxicology papers focus on single endpoints. Full batteries are rarely reported^7^. Reports describing coordinated evaluation of structural, functional, and cellular immune parameters within a single integrated experimental framework remain comparatively limited. In addition, many experimental investigations are not conducted under Good Laboratory Practice (GLP) conditions, which may reduce their direct applicability in regulatory decision-making contexts^8^. Demonstration of concordance among multiple immunotoxicity endpoints generated within a controlled study design may therefore improve interpretability of immune findings and contribute to more transparent hazard characterization within regulatory risk assessment.

Cyclophosphamide is a well-characterized cytotoxic alkylating agent with established immunosuppressive activity resulting primarily from its effects on proliferating lymphocytes. Its ability to suppress both humoral and cellular immune responses has been extensively documented in experimental immunotoxicology and it has frequently been used as a reference compound in immune function assays^5^. Owing to its consistent and dose-dependent immunosuppressive effects, cyclophosphamide represents an appropriate model compound for demonstrating application of integrated immunotoxicity assessment strategies.

The present study was therefore designed to illustrate implementation of an integrated immunotoxicity evaluation consistent with regulatory principles of hazard identification and weight-of-evidence interpretation. Using cyclophosphamide monohydrate in male Sprague–Dawley rats, investigations were conducted under Good Laboratory Practice (GLP) conditions to examine coordinated endpoints including clinical pathology indicators of immune status, lymphoid organ assessment, functional immune assays, and cellular immunophenotyping. Evaluation of these complementary datasets within a single experimental framework was intended to provide a practical example of integrated hazard characterization and interpretation of immunotoxic potential within a regulatory toxicology context.

## Materials and Methods

### Study Design and Animal Husbandry

Male Sprague–Dawley rats (12 weeks old) were obtained from RCC Laboratories India Pvt. Ltd., a facility approved by the Committee for Control and Supervision of Experiments on Animals (CCSEA) and accredited by the Association for Assessment and Accreditation of Laboratory Animal Care International (AAALAC). Animals were housed under controlled environmental conditions (temperature 19–24 °C, relative humidity 59–68%, 12 h light/dark cycle) with ad libitum access to standard laboratory diet (AF-2000 SP, Krishna Valley Agrotech LLP) and UV-purified water.

The study was approved by the Institutional Animal Ethics Committee and conducted in accordance with OECD Good Laboratory Practice (GLP) principles. Animals were randomized by body-weight stratification into three groups (n = 6 per group): vehicle control, cyclophosphamide 50 mg/kg, and cyclophosphamide 100 mg/kg.

### Test Item and Dose Administration

Cyclophosphamide monohydrate (Sigma-Aldrich; purity 99.4%) was dissolved in sterile 0.9% sodium chloride solution to obtain dosing formulations of 5 mg/mL and 10 mg/mL. Animals received intraperitoneal injections on study days 1 and 4 at a dose volume of 10 mL/kg. The dosing regimen was selected based on published immunotoxicity models designed to induce lymphoid suppression while maintaining animal viability^9^.

### Immunization with Sheep Red Blood Cells

Sheep red blood cells (SRBC) were used as the antigen for evaluation of humoral and cellular immune responses. Sheep blood collected in Alsever’s solution was washed with physiological saline and adjusted to the required concentrations. For induction of the antibody response, animals received an intraperitoneal injection of 1 mL of 10% SRBC suspension four days prior to necropsy. For delayed-type hypersensitivity (DTH) evaluation, 100 µL of 1% SRBC suspension was injected into the right hind footpad.

### Clinical Observations and Body Weight

Animals were observed daily for mortality, clinical signs, and general health. Body weights were recorded during acclimatization and at regular intervals throughout the dosing period.

### Clinical Pathology Haematology

On study day 8, blood samples were collected from the retro-orbital plexus under anaesthesia into K₂-EDTA tubes. Haematological parameters including erythrocyte count, haemoglobin, haematocrit, red cell indices (MCV, MCH, MCHC), platelet count, total leukocyte count, and differential leukocyte count were measured using an automated haematology analyser (Sysmex XN-330).

### Clinical Biochemistry

Serum samples obtained from blood collected in plain tubes were analyzed using a Cobas 111 clinical chemistry analyzer (Roche) for total protein, albumin, and globulin concentrations. Albumin–globulin ratio was calculated.

### Humoral Immune Response (Hemagglutination Assay)

Serum antibody response to SRBC was determined using a hemagglutination (HA) assay. Serum samples collected prior to treatment and four days after immunization were serially diluted two-fold in microtiter plates. Equal volumes of 1% SRBC suspension were added to each well and plates were incubated at 37 °C followed by overnight refrigeration. The reciprocal of the highest serum dilution showing visible agglutination was recorded as the antibody titer and expressed as log₂ values.

### Delayed-Type Hypersensitivity (DTH) Response

Cell-mediated immune response was evaluated using the SRBC-induced DTH model. On day 7, 100 µL of 1% SRBC suspension was injected into the right hind footpad, while the left hind footpad received saline as control. Footpad thickness was measured using a digital caliper prior to challenge and at 6 h and 24 h post-injection. The DTH response was expressed as the increase in footpad thickness relative to baseline.

### Immunophenotyping (TBNK Analysis)

Peripheral blood immunophenotyping was performed using flow cytometry (BD FACSVerse™). Whole blood (100 µL) was incubated with fluorochrome-conjugated monoclonal antibodies specific for rat leukocyte markers to identify T cells, B cells, and natural killer (NK) cells (**Table 1**). After red blood cell lysis and washing, samples were analyzed by flow cytometry and approximately 10,000 lymphocyte events were acquired per sample. Results were expressed as percentages of lymphocyte subsets and used for calculation of absolute cell counts.

**Table 1.**
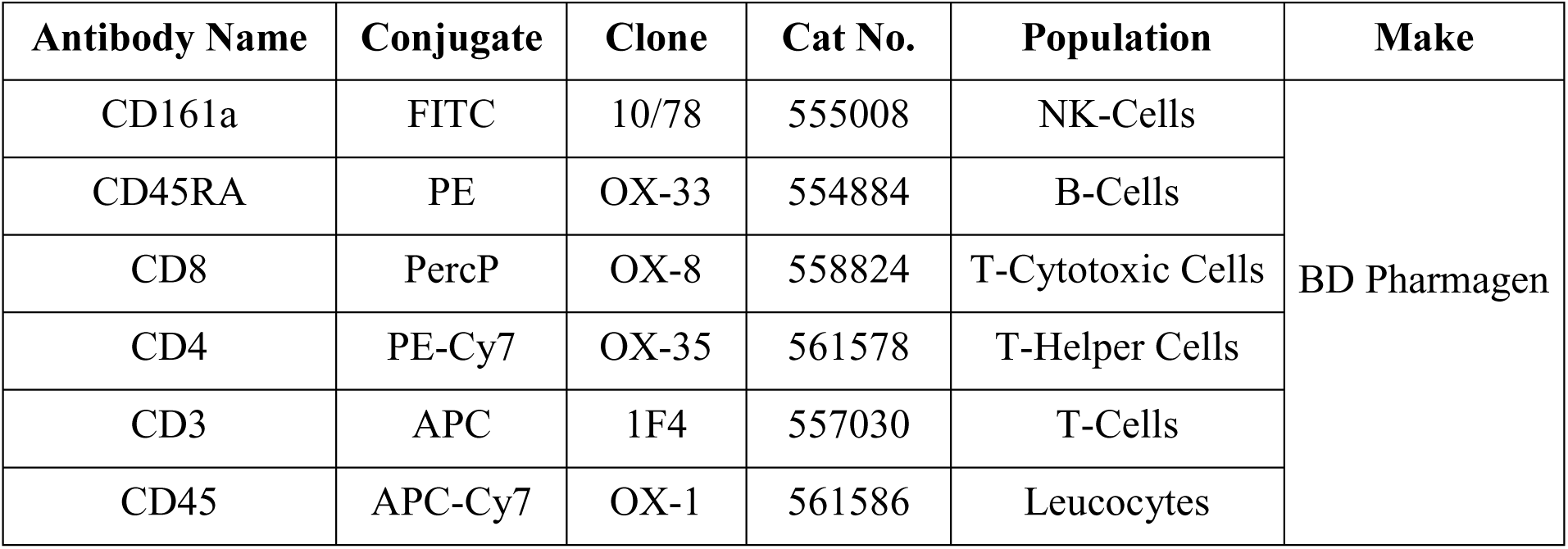
Monoclonal Antibodies Used for Rat TBNK Analysis.

### Necropsy, Organ Weights and Histopathology

Animals were euthanized on study day 8 by CO₂ inhalation followed by exsanguination. A complete necropsy was performed and organs including spleen, thymus, liver, and kidneys were weighed. Tissues including spleen, thymus, lymph nodes, liver, kidney, ileum with Peyer’s patches, and sternum (bone marrow) were fixed in 10% neutral buffered formalin. Formalin-fixed tissues were processed using standard histological techniques, embedded in paraffin, sectioned at 4–5 µm, stained with hematoxylin and eosin, and examined microscopically. Bone marrow smears were stained with May–Grünwald–Giemsa for differential cell counts and evaluation of myeloid-to-erythroid ratio.

### Statistical Analysis

Data were analyzed using StatPlus software. Continuous variables were expressed as mean ± standard deviation (SD). Normality was assessed using the Shapiro–Wilk test. Parametric data were analyzed using one-way analysis of variance (ANOVA) followed by Fisher’s least significant difference test for comparison between groups. Non-parametric data were analyzed using Kruskal–Wallis analysis with appropriate post-hoc comparisons. Statistical significance was considered at *p* ≤ 0.05.

## Results

### Hazard Identification

#### Clinical Observations

No mortality occurred during the acclimatization or treatment period. Cage-side observations did not reveal any treatment-related clinical signs in control or cyclophosphamide-treated animals throughout the study (**Table 2)**.

**Table 2.** Summary of Mortality and Observed Clinical Signs in Control and Cyclophosphamide-Treated Groups rats.

| Parameters | Control | Cyclophosphamide<br>50 mg/kg body weight | Cyclophosphamide<br>100 mg/kg body weight |
| --- | --- | --- | --- |
| Mortality | Nil | Nil | Nil |
| Clinical Signs | Nil | Nil | Nil |

Control animals exhibited a gradual increase in body weight during the study period (**Table 3)**. In contrast, animals receiving cyclophosphamide showed reduced body weight gain relative to baseline values, with a more evident downward trend at the high dose (100 mg/kg bw). Although absolute body weights did not differ significantly between groups, analysis of relative body weight indicated suppression of weight gain in treated animals (**Table 4)**. A statistically significant reduction in relative body weight was observed in the low-dose group on Day 6 when compared with controls. These observations suggest mild systemic effects associated with cyclophosphamide administration but without overt clinical toxicity.

**Table 3.** Mean Absolute Body Weight (g) of Control and Cyclophosphamide-Treated Rats.

| Groups | Day 1 | Day 3 | Day 6 | Day 8 |
| --- | --- | --- | --- | --- |
| Control | 333.04 ±<br>25.78 | 338.89 ±<br>24.72 | 346.70 ±<br>25.97 | 338.28 ±<br>28.46 |
| Cyclophosphamide<br>50 mg/kg body weight | 346.48 ±<br>25.49 | 325.10 ±<br>20.51 | 323.25 ±<br>23.22 | 325.41 ±<br>17.44 |
| Cyclophosphamide<br>100 mg/kg body<br>weight | 325.44 ±<br>23.59 | 316.69 ±<br>20.37 | 319.23 ±<br>22.78 | 313.00 ±<br>22.42 |

Data are expressed as mean ± standard deviation (n = 6 per group). Differences were considered statistically significant at $p \leq 0.05$ . Means with the same superscript letter do not differ significantly between groups.

**Table 4.** Relative Body Weight (%) of Control and Cyclophosphamide-Treated Rats During the Study Period.

| **Group** | **Day 3** | **Day 6** | **Day 8** |
| --- | --- | --- | --- |
| **Control** | 100.79<br>±<br>1.16 | 106.13<br>±<br>1.52 | 101.54<br>±<br>2.17 |
| **Cyclophosphamide @ 50 mg/Kg Body weight** | 94.33<br>±<br>9.59 | 93.65<sup>a</sup><br>±<br>8.63 | 94.41<br>±<br>9.02 |
| **Cyclophosphamide @ 100 mg/Kg Body weight** | 97.36<br>±<br>1.21 | 98.10<br>±<br>1.01 | 96.19<br>±<br>1.46 |

Data are expressed as mean ± standard deviation (n = 6 per group). Differences were considered statistically significant at $p \leq 0.05$ . Means with the same superscript letter do not differ significantly between groups.

#### Hematology

Cyclophosphamide treatment produced dose-related alterations in hematological parameters (**Table 5)**. Red blood cell count, hemoglobin concentration, and hematocrit values showed decreases in treated animals, with greater reductions observed at the high dose. Mean corpuscular indices (MCV, MCH, and MCHC) remained comparable to control values.

**Table 5.**
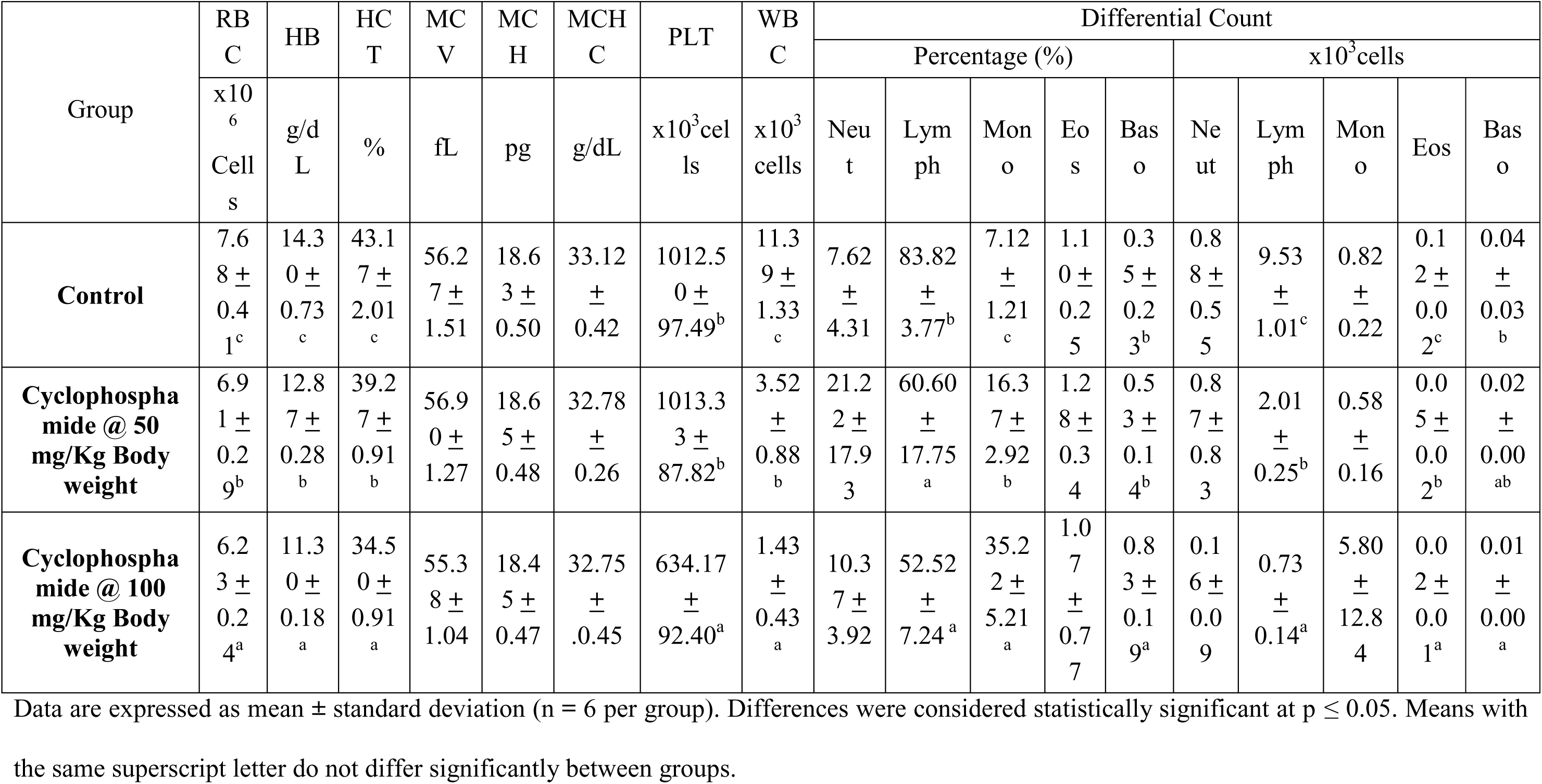
Summary of Hematological Parameters in Control and Cyclophosphamide-Treated Rats.

Total white blood cell counts were reduced in treated groups, reflecting a marked decline in lymphocyte numbers. Differential leukocyte counts showed relative neutrophilia and monocytosis, accompanied by a pronounced reduction in lymphocyte counts, particularly in animals receiving 100 mg/kg bw. Platelet counts were largely unaffected at the low dose but showed a reduction at the high dose.

These findings indicate suppression of circulating leukocyte populations, consistent with the known myelosuppressive activity of cyclophosphamide and providing early evidence of immune system involvement.

#### Clinical Biochemistry

Details of serum total protein, albumin, globulin, and albumin-to-globulin (A:G) ratio are presented in Table 6. Cyclophosphamide administration resulted in a statistically significant, dose-dependent reduction in serum total protein levels in both treated groups, with a more pronounced decrease at the high dose. Serum albumin concentrations remained comparable across all groups. Calculated globulin levels (total protein minus albumin) showed a dose-dependent reduction, reaching statistical significance at the high dose. As a consequence of the selective reduction in the globulin fraction, the A:G ratio increased progressively with dose, although the change did not reach statistical significance.

**Table 6.** Summary of Clinical Biochemistry Parameters in Control and Cyclophosphamide-Treated Rats.

| Group | Total protien (g/L) | Albumin (g/L ) | Globulin (g/L) | A/G Ratio |
| --- | --- | --- | --- | --- |
| Control | 59.88 ± 2.30 | 41.42 ± 1.45 | 18.45 ± 1.38 <sup>b</sup> | 2.23 ± 0.16 |
| Cyclophosphamide @ 50 mg/Kg Body weight | 57.30 ± 1.44 <sup>a</sup> | 40.00 ± 1.88 | 17.28 ± 2.42 <sup>ab</sup> | 2.37 ± 0.44 |
| Cyclophosphamide @ 100 mg/Kg Body weight | 55.80 ± 1.52 <sup>a</sup> | 40.22 ± 1.22 | 15.57 ± 1.08 <sup>a</sup> | 2.62 ± 0.19 |

Data are expressed as mean ± standard deviation (n = 6 per group). Differences were considered statistically significant at $p \leq 0.05$ . Means with the same superscript letter do not differ significantly between groups.

### Necropsy, Organ Weights and Histopathology

Evaluation of organ weights demonstrated selective effects on lymphoid organs (**Table 7 and 8**). Absolute and relative weights of spleen and thymus were significantly reduced in both treatment groups, with greater reductions observed at 100 mg/kg bw. In contrast, liver and kidney weights remained comparable to those of control animals.

**Table 7.** Absolute Organ Weights in Control and Cyclophosphamide-Treated Rats.

| Group | Kidney | Liver | Spleen | Thymus |
| --- | --- | --- | --- | --- |
| Control | 2.30 ± 0.20 | 11.91 ± 1.49 | 0.49 ± 0.09<sup>b</sup> | 0.89 ± 0.09<sup>b</sup> |
| Cyclophosphamide @ 50 mg/Kg Body weight | 2.31 ± 0.12 | 12.77 ± 1.17 | 0.35 ± 0.08<sup>a</sup> | 0.64 ± 0.08<sup>a</sup> |
| Cyclophosphamide @ 100 mg/Kg Body weight | 2.35 ± 0.26 | 12.49 ± 1.41 | 0.28 ± 0.08<sup>a</sup> | 0.54 ± 0.11<sup>a</sup> |

Data are expressed as mean ± standard deviation (n = 6 per group). Differences were considered statistically significant at $p \leq 0.05$ . Means with the same superscript letter do not differ significantly between groups.

**Table 8.** Relative Organ Weights in Control and Cyclophosphamide-Treated Rats.

| **Group** | **Kidney** | **Liver** | **Spleen** | **Thymus** |
| --- | --- | --- | --- | --- |
| **Control** | 0.68 ± 0.04 | 3.52 ± 0.22 | 0.15 ± 0.03<sup>b</sup> | 0.26 ± 0.02<sup>b</sup> |
| **Cyclophosphamide**<br>**@ 50 mg/Kg Body**<br>**weight** | 0.71 ± 0.05 | 3.94 ± 0.48 | 0.11 ± 0.03<sup>a</sup> | 0.20 ± 0.03<sup>b</sup> |
| **Cyclophosphamide**<br>**@ 100 mg/Kg Body**<br>**weight** | 0.75 ± 0.06 | 3.99 ± 0.40 | 0.09 ± 0.03<sup>a</sup> | 0.17 ± 0.03<sup>a</sup> |

Data are expressed as mean ± standard deviation (n = 6 per group). Differences were considered statistically significant at $p \leq 0.05$ . Means with the same superscript letter do not differ significantly between groups.

The decrease in spleen and thymus weights indicates structural effects on primary and secondary lymphoid organs, consistent with suppression of lymphoid cell populations.

Macroscopic examination at necropsy did not reveal abnormalities in liver or kidney tissues. However, spleen and thymus from treated animals appeared smaller and paler compared with controls, particularly at the high dose **(Table 9)**.

**Table 9.** Summary of Macroscopic Findings in Control and Cyclophosphamide-Treated Rats (n/N)

| **Macroscopic Findings** | **Control** | **Cyclophosphamide**<br>**@ 50 mg/Kg Body**<br>**weight** | **Cyclophosphamide**<br>**@ 100 mg/Kg Body**<br>**weight** |
| --- | --- | --- | --- |
| **EXTERNAL** |  |  |  |
| No abnormality detected | 6/6 | 6/6 | 6/6 |
| **INTERNAL** |  |  |  |
| No abnormality detected | 6/6 | 1/6 | 0/6 |
| Spleen reduced in size | - | 1/6 | 3/6 |
| Thymus reduced in size | - | 5/6 | 5/6 |

(Where n = number of animals showing lesion, N = total number of animals per group)

Microscopic examination confirmed treatment-related alterations in lymphoid tissues **(Table 10**; **Figure 1)**. In the spleen, lymphoid depletion and atrophy were observed in treated animals, with severity ranging from mild to severe. Thymic sections demonstrated lymphocyte depletion and cortical atrophy in most animals from the treatment groups.

**Figure 1.**
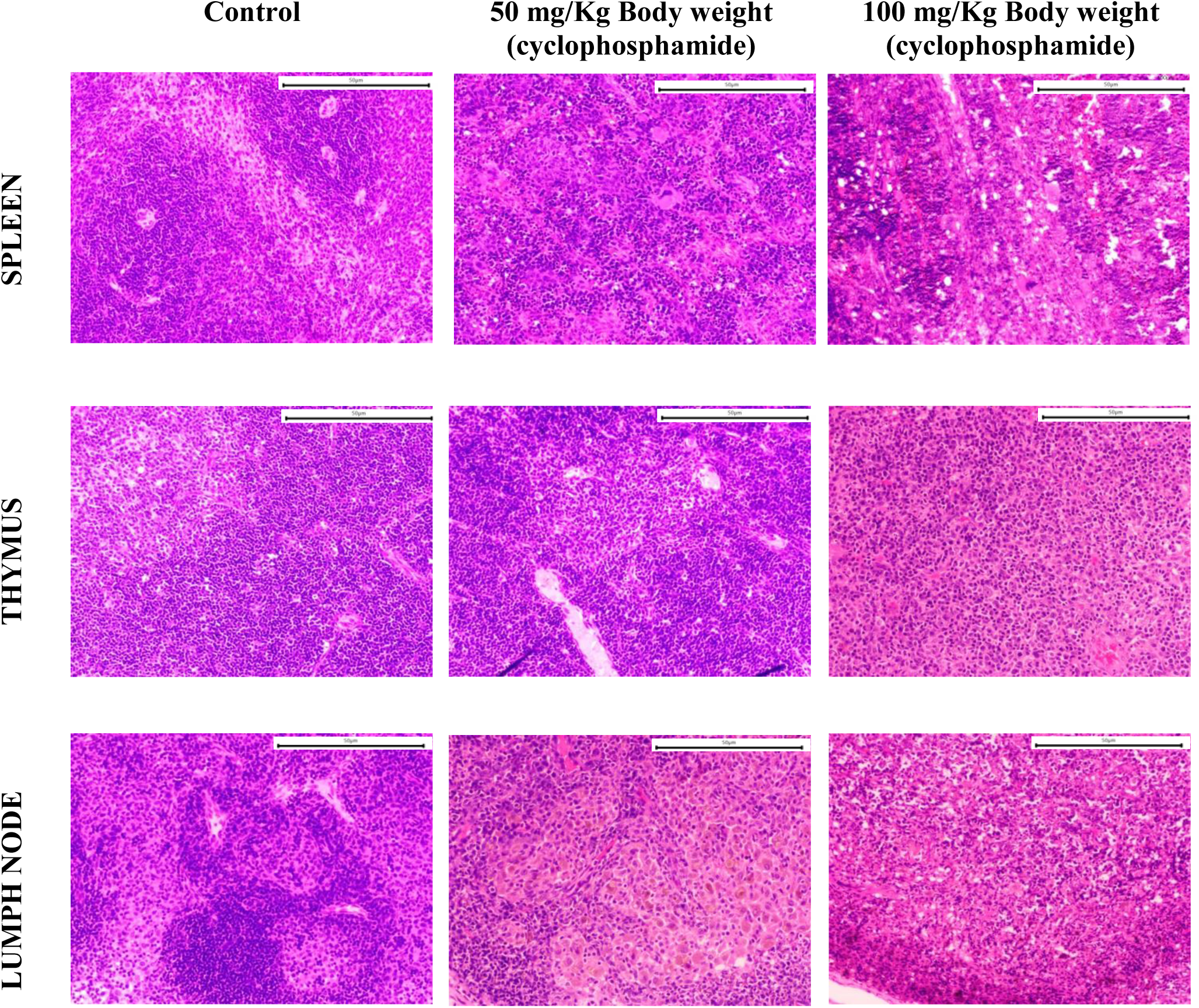
Histopathological changes in lymphoid organs following cyclophosphamide exposure. Hematoxylin and eosin (H&E)-stained sections of spleen and thymus from control and treated groups. Control animals exhibited normal follicular architecture with intact germinal centers. Cyclophosphamide-treated animals showed dose-dependent lymphoid depletion, cortical thinning in thymus, and reduced white pulp cellularity in spleen. Magnification: 200×. Scale bar: 50 μm.

**Table 10.**
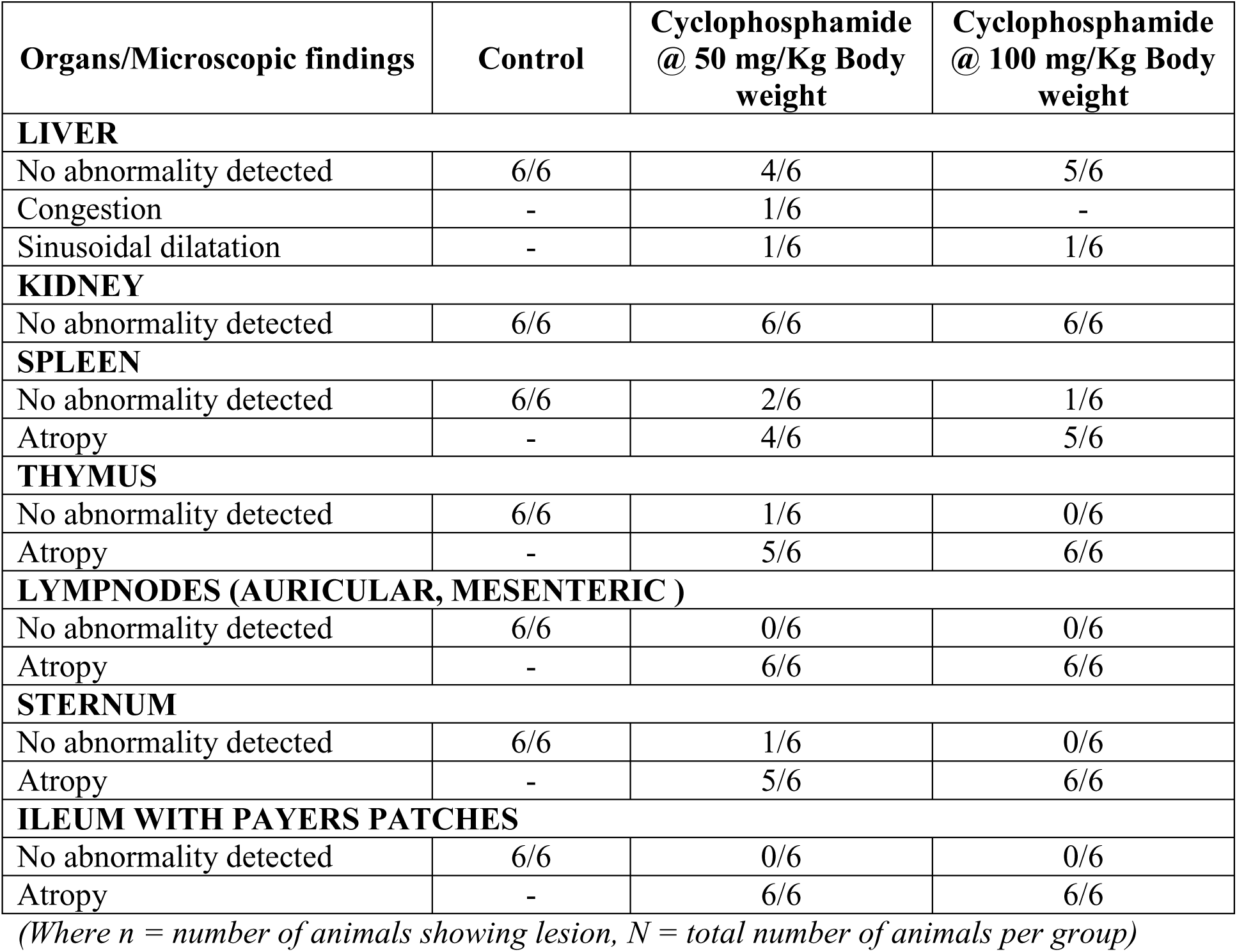
Microscopic Findings in Control and Cyclophosphamide-Treated Rats (n/N)

| **Organs/Microscopic findings** | **Control** | **Cyclophosphamide**<br>**@ 50 mg/Kg Body**<br>**weight** | **Cyclophosphamide**<br>**@ 100 mg/Kg Body**<br>**weight** |
| --- | --- | --- | --- |
| **LIVER** |  |  |  |
| No abnormality detected | 6/6 | 4/6 | 5/6 |
| Congestion | - | 1/6 | - |
| Sinusoidal dilatation | - | 1/6 | 1/6 |
| **KIDNEY** |  |  |  |
| No abnormality detected | 6/6 | 6/6 | 6/6 |
| **SPLEEN** |  |  |  |
| No abnormality detected | 6/6 | 2/6 | 1/6 |
| Atropy | - | 4/6 | 5/6 |
| **THYMUS** |  |  |  |
| No abnormality detected | 6/6 | 1/6 | 0/6 |
| Atropy | - | 5/6 | 6/6 |
| **LYMPNODES (AURICULAR, MESENTERIC )** |  |  |  |
| No abnormality detected | 6/6 | 0/6 | 0/6 |
| Atropy | - | 6/6 | 6/6 |
| **STERNUM** |  |  |  |
| No abnormality detected | 6/6 | 1/6 | 0/6 |
| Atropy | - | 5/6 | 6/6 |
| **ILEUM WITH PAYERS PATCHES** |  |  |  |
| No abnormality detected | 6/6 | 0/6 | 0/6 |
| Atropy | - | 6/6 | 6/6 |

*(Where n = number of animals showing lesion, N = total number of animals per group)*

Similar findings were noted in other lymphoid compartments. Lymph nodes from treated animals exhibited mild to moderate atrophy. Bone marrow examination revealed reduced cellularity and alterations in lineage distribution, with decreases in myeloid progenitor populations (**Table 11**, **Figure 2)**. Peyer’s patches in the ileum also showed lymphoid depletion in treated animals.

**Figure 2.**
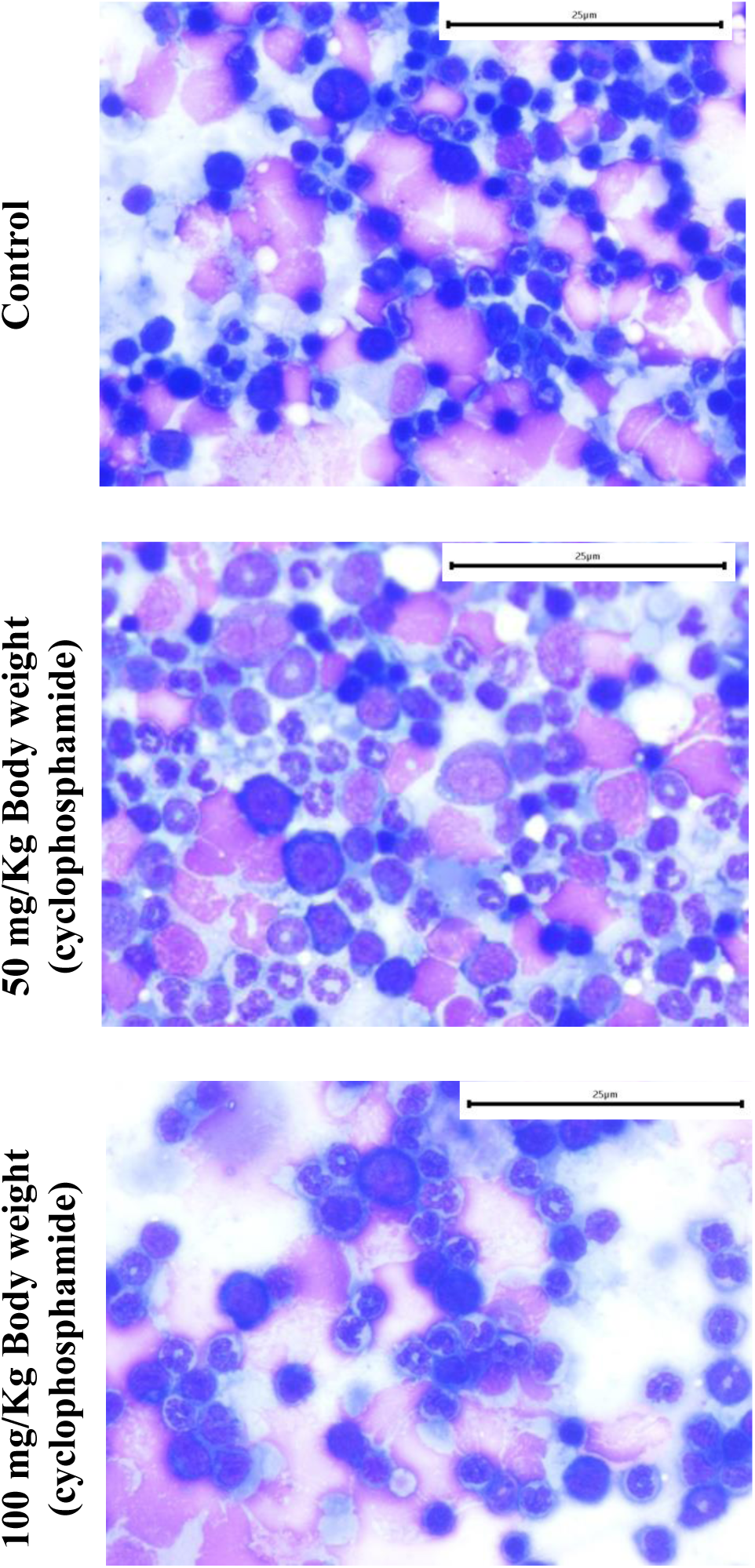
Bone marrow cytology following cyclophosphamide administration. Representative May-Grünwald-Giemsa–stained smears from sternum bone marrow. Control animals showed balanced myeloid and erythroid populations. Cyclophosphamide-treated dose animals exhibited hypocellularity, reduced erythroid precursors, and altered M:E ratios consistent with myelosuppression. Magnification: 400×. Scale bar: 25 μm.

**Table 11.** Bone Marrow Differential Count and Myeloid:Erythroid (M:E) Ratio in Control and Cyclophosphamide-Treated Rats.

| Group | Myeloid cells | Erythroid cells | M:E ratio |
| --- | --- | --- | --- |
| Control | 97.00 ± 10.32<sup>b</sup> | 103.00 ± 10.32<sup>b</sup> | 0.96 ± 0.20<sup>b</sup> |
| Cyclophosphamide @ 50 mg/Kg Body weight | 87.50 ± 17.11<sup>b</sup> | 112.50 ± 17.11<sup>b</sup> | 0.82 ± 0.31<sup>b</sup> |
| Cyclophosphamide @ 100 mg/Kg Body weight | 62.17 ± 14.91<sup>a</sup> | 137.83 ± 14.91<sup>a</sup> | 0.47 ± 0.16<sup>a</sup> |

Note- bone marrow differential counts (200 nucleated cells/animal): myeloid cells, erythroid cells, and myeloid-to-erythroid (M:E) ratio Data are expressed as mean ± standard deviation (n = 6 per group). Differences were considered statistically significant at $p \leq 0.05$ . Means with the same superscript letter do not differ significantly between groups.

These histopathological findings demonstrate consistent structural suppression across multiple lymphoid tissues and hematopoietic compartments.

#### Functional Validation

To determine if the observed structural atrophy in lymphoid tissues correlated with functional impairment, we evaluated adaptive immune competence through TDAR and DTH assays.

#### Hemagglutination Antibody Response

Baseline hemagglutination antibody titers measured prior to treatment were comparable across groups, indicating similar pre-immunization immune status.

Following immunization with sheep red blood cells, control animals developed strong antibody responses, as reflected by high hemagglutination titers. In contrast, animals treated with cyclophosphamide showed marked suppression of antibody production. Both dose groups exhibited significantly lower antibody titers compared with controls, with a clear reduction in humoral immune response (**Table 12**).

**Table 12.** Hemagglutination Antibody Titer in Control and Cyclophosphamide-Treated Rats.

| Group | Antibody titer* |  | Log 2 antibody titer |  |
| --- | --- | --- | --- | --- |
|  | Day 1 | Day 8 | Day 1 | Day 8 |
| Control | 2.67 ± 1.03 | 173.33 ± 99.36<sup>b</sup> | 1.33 ± 0.52 | 7.00 ± 1.55<sup>b</sup> |
| Cyclophosphamide @ 50 mg/Kg Body weight | 3.00 ± 2.76 | 19.33 ± 14.79<sup>a</sup> | 1.33 ± 1.03 | 3.50 ± 1.97<sup>a</sup> |
| Cyclophosphamide @ 100 mg/Kg Body weight | 1.00 ± 1.10 | 6.00 ± 2.19<sup>a</sup> | 0.50 ± 0.55 | 2.50 ± 0.55<sup>a</sup> |

Data are expressed as mean ± standard deviation (n = 6 per group). Differences were considered statistically significant at $p \leq 0.05$ . Means with the same superscript letter do not differ significantly between groups.

These results demonstrate that cyclophosphamide treatment impairs the T-dependent antibody response, indicating functional suppression of adaptive humoral immunity.

#### Delayed-Type Hypersensitivity (DTH)

Evaluation of delayed-type hypersensitivity responses revealed minimal differences among groups at 6 hours after antigen challenge. However, clear differences were evident at the 24-hour measurement (**Table 13**).

**Table 13.** Delayed-Type Hypersensitivity (DTH) Paw Edema Response Following SRBC Challenge: Absolute and Percentage Changes in Left and Right Paw Thickness at 6 and 24 Hours in Rats.

| Group | Left Paw |  |  |  | Right Paw |  |  |  |
| --- | --- | --- | --- | --- | --- | --- | --- | --- |
|  | Absolute Difference in paw after challenge (mm) |  | % Change in paw after challenge |  | Absolute Difference in paw after challenge (mm) |  | % Change in paw after challenge |  |
|  | 6hr | 24hr | 6hr | 24hr | 6hr | 24hr | 6hr | 24hr |
| Control | 0.03 ± 0.08 | 0.05 ± 0.08 | 1.00 ± 2.27 | 1.55 ± 2.35 | 0.13 ± 0.28 | 1.47 ± 0.72 | 4.14 ± 8.90 | 43.37 ± 22.72 |
| Cyclophosphamide @ 50 mg/Kg Body weight | 0.05 ± 0.19 | 0.04 ± 0.12 | 1.45 ± 5.67 | 1.03 ± 3.71 | 0.03 ± 0.08 | 0.09 ± 0.11<sup>a</sup> | 0.78 ± 2.31 | 2.72 ± 3.32<sup>a</sup> |
| Cyclophosphamide @ 100 mg/Kg Body weight | 0.05 ± 0.07 | 0.05 ± 0.02 | 1.54 ± 2.13 | 1.38 ± 0.67 | 0.02 ± 0.14 | 0.03 ± 0.05<sup>a</sup> | 0.59 ± 4.13 | 0.92 ± 1.47<sup>a</sup> |

Data are expressed as mean ± standard deviation (n = 6 per group). Differences were considered statistically significant at $p \leq 0.05$ . Means with the same superscript letter do not differ significantly between groups.

Control animals exhibited substantial paw swelling in the antigen-challenged paw, indicating a robust cell-mediated immune response. In contrast, both cyclophosphamide-treated groups showed markedly reduced paw swelling. Suppression of the DTH response was statistically significant compared with controls. A dose-related difference between treatment groups was observed for absolute paw thickness change.

No significant changes were observed in saline-injected paws, confirming the antigen specificity of the response. These findings indicate impairment of T-cell–mediated immune responses following cyclophosphamide exposure.

#### Cellular Characterization (Lymphocyte Subset Alterations)

Flow cytometric immunophenotyping was performed to confirm involvement of both CD3⁺ T lymphocytes and CD45RA⁺ B lymphocytes in treated cohorts (**Figure 3**)

**Figure 3.**
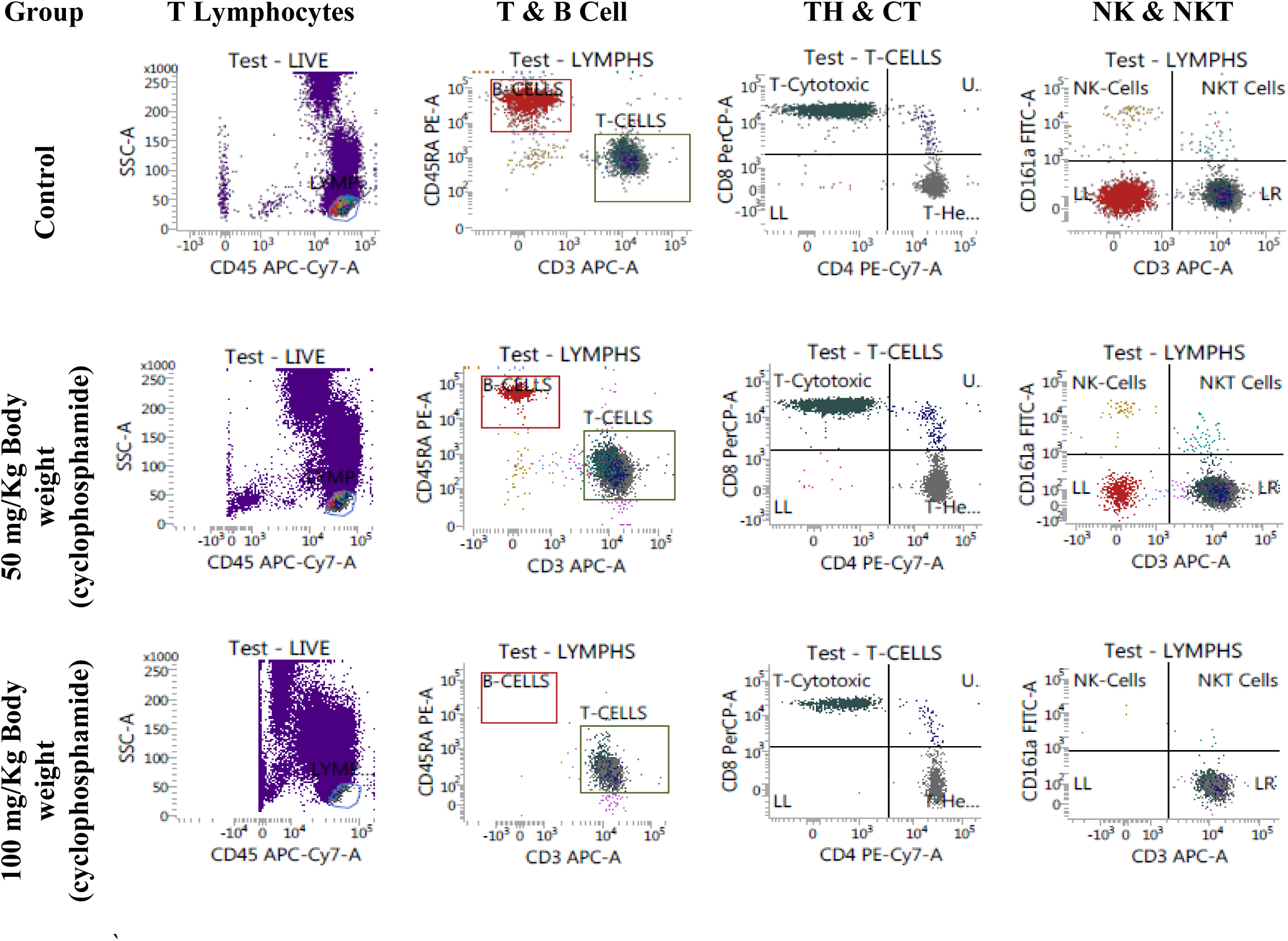
Flow cytometry analysis of lymphocyte subsets in peripheral blood. Representative dot plots showing CD45^+^ Leukocytes, CD45RA⁺ B cells, CD3⁺ T cells, CD4^+^ T-helper cells, CD8^+^ T-cytotoxic cells and CD161a⁺ NK cells in control, low-dose (50 mg/kg), and high-dose (100 mg/kg) cyclophosphamide-treated rats on Day 8. Cyclophosphamide induced dose-dependent lymphopenia, with marked depletion of B cells and reduced CD4⁺ T-helper populations. Data are expressed as mean absolute counts ± SD (n=6/group)

#### Lymphocyte Immunophenotyping (TBNK Analysis)

Flow cytometric analysis revealed significant treatment-related changes in lymphocyte populations. Total lymphocyte percentages (**Table 14)** and absolute counts (**Table 15)** decreased markedly in cyclophosphamide-treated animals, with pronounced lymphopenia observed at the high dose. Although the relative proportion of CD3⁺ T cells increased, absolute T-cell numbers declined, particularly in the high-dose group. B-cell populations showed the greatest sensitivity to treatment, with substantial reductions in both percentage and absolute cell counts. Near-complete depletion of B cells was observed in animals receiving 100 mg/kg bw.

**Table 14.**
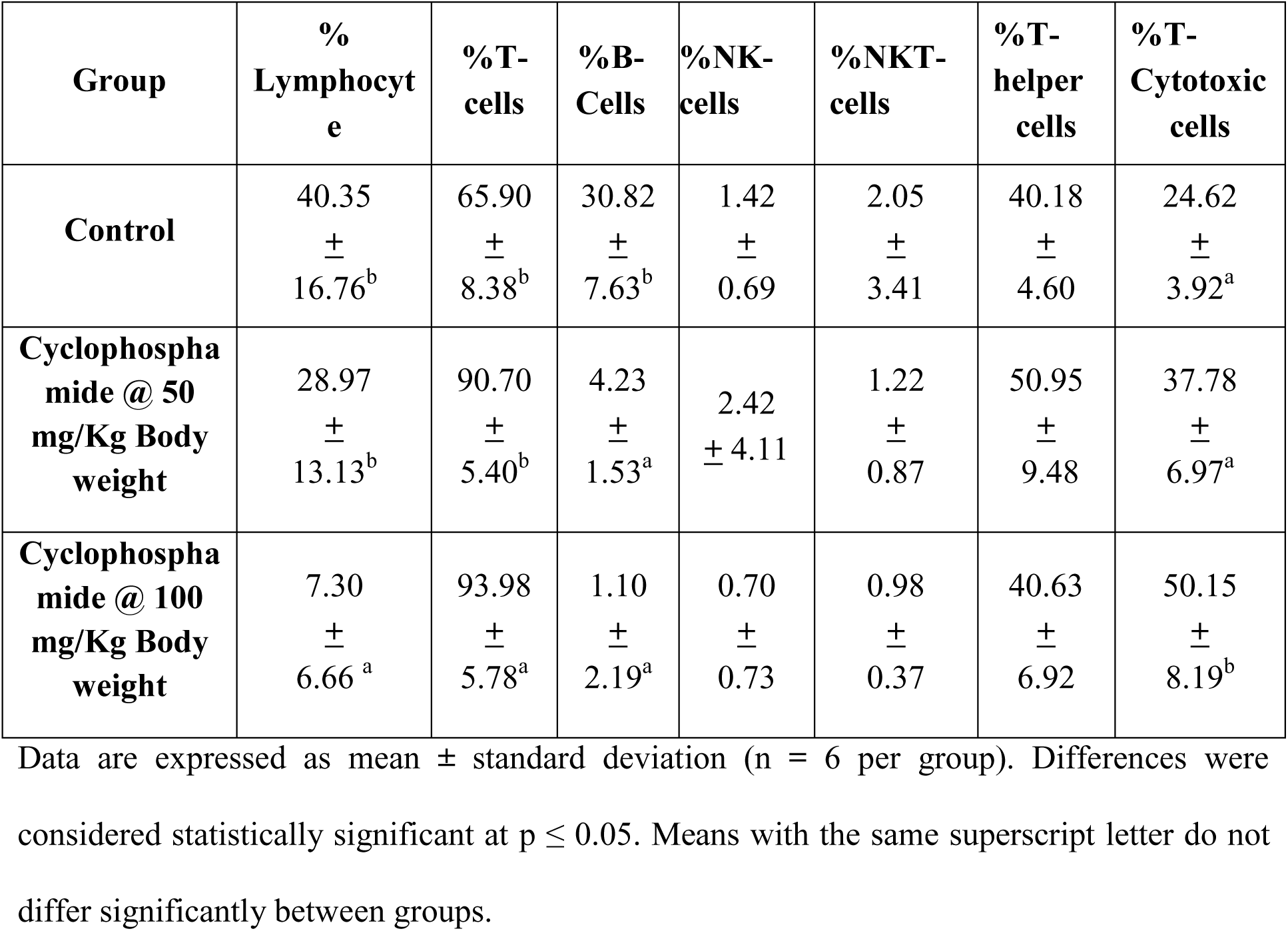
Immunophenotyping of Lymphocyte Subpopulations (% Cells) in Control and Cyclophosphamide-Treated Rats.

| **Group** | **%**<br>**Lymphocyte** | **%T-**<br>**cells** | **%B-**<br>**Cells** | **%NK-**<br>**cells** | **%NKT-**<br>**cells** | **%T-**<br>**helper**<br>**cells** | **%T-**<br>**Cytotoxic**<br>**cells** |
| --- | --- | --- | --- | --- | --- | --- | --- |
| **Control** | 40.35<br>±<br>16.76<sup>b</sup> | 65.90<br>±<br>8.38<sup>b</sup> | 30.82<br>±<br>7.63<sup>b</sup> | 1.42<br>±<br>0.69 | 2.05<br>±<br>3.41 | 40.18<br>±<br>4.60 | 24.62<br>±<br>3.92<sup>a</sup> |
| **Cyclophosphamide @ 50**<br>**mg/Kg Body**<br>**weight** | 28.97<br>±<br>13.13<sup>b</sup> | 90.70<br>±<br>5.40<sup>b</sup> | 4.23<br>±<br>1.53<sup>a</sup> | 2.42<br>± 4.11 | 1.22<br>±<br>0.87 | 50.95<br>±<br>9.48 | 37.78<br>±<br>6.97<sup>a</sup> |
| **Cyclophosphamide @ 100**<br>**mg/Kg Body**<br>**weight** | 7.30<br>±<br>6.66<sup>a</sup> | 93.98<br>±<br>5.78<sup>a</sup> | 1.10<br>±<br>2.19<sup>a</sup> | 0.70<br>±<br>0.73 | 0.98<br>±<br>0.37 | 40.63<br>±<br>6.92 | 50.15<br>±<br>8.19<sup>b</sup> |

Data are expressed as mean ± standard deviation (n = 6 per group). Differences were considered statistically significant at $p \leq 0.05$ . Means with the same superscript letter do not differ significantly between groups.

**Table 15.** Absolute Counts of Lymphocyte Subpopulations in Control and Cyclophosphamide-Treated Rats.

| **Group** | **Lymphocytes** | **T-cells** | **B-cells** | **NK-cells** | **NKT-cells** | **T-helper cells** | **T-Cytotoxic cells** |
| --- | --- | --- | --- | --- | --- | --- | --- |
| **Control** | 4035.00<br>±<br>1676.46<sup>b</sup> | 2567.7<br>5<br>±<br>995.91<sub>b</sub> | 1335.<br>90<br>±<br>701.1<br>1<sup>b</sup> | 59.45<br>±<br>43.15 | 39.63<br>±<br>22.96 | 1014.0<br>7<br>±<br>398.35<sup>b</sup> | 607.71<br>±<br>221.87<sup>b</sup> |
| **Cyclophosphamide @ 50 mg/Kg Body weight** | 2896.67<br>±<br>1313.48<sup>b</sup> | 2594.6<br>4<br>±<br>1109.2<br>0<sup>b</sup> | 118.3<br>3<br>±<br>66.65<sup>a</sup> | 97.98<br>±<br>182.07 | 43.65<br>±<br>43.06 | 1252.2<br>3<br>±<br>399.39<sup>b</sup> | 1028.36<br>±<br>580.38<sup>b</sup> |
| **Cyclophosphamide @ 100 mg/Kg Body weight** | 730.00<br>±<br>665.67<sup>a</sup> | 657.70<br>±<br>554.36<sub>a</sub> | 18.97<br>±<br>40.34<sup>a</sup> | 8.73<br>±<br>12.83 | 8.63<br>±<br>9.69 | 279.33<br>±<br>248.65<sup>a</sup> | 298.48<br>±<br>218.77<sup>a</sup> |

Data are expressed as mean ± standard deviation (n = 6 per group). Differences were considered statistically significant at $p \leq 0.05$ . Means with the same superscript letter do not differ significantly between groups.

Both CD4⁺ helper and CD8⁺ cytotoxic T-cell subsets exhibited reductions in absolute counts at the high dose. In contrast, natural killer (NK) and NKT cell populations showed minimal changes. These immunophenotyping data demonstrate selective depletion of adaptive immune cell populations, particularly B lymphocytes, providing mechanistic evidence supporting the observed suppression of humoral and cellular immune responses.

## Discussion

### Hazard Characterization: Structural Immunotoxicity Indicators

Cyclophosphamide administration produced reductions in spleen and thymus weights alongside histopathological evidence of lymphoid depletion. Lymphoid atrophy occurred in the spleen, thymus, lymph nodes, Peyer’s patches, and bone marrow, affecting both primary and secondary immune organs. These changes align with effects of cytotoxic agents that impair lymphocyte proliferation and survival (10). Concordance between organ weight reductions and histopathological lymphocyte depletion confirms morphological immune suppression. Early hematological alterations, including reduced total leukocyte counts driven by lymphocyte depletion, further indicated hematopoietic suppression, with bone marrow showing decreased myeloid progenitors and myeloid-to-erythroid ratio (11,12).

### Confirmation of Immune Impairment: Functional Assay Analysis

Functional assays demonstrated marked immune incompetence. Hemagglutination antibody responses to SRBC immunization were strongly suppressed in treated animals compared to controls (13). This T-dependent antibody response requires coordinated antigen-presenting cells, T-helper cells, and B-lymphocytes, marking it as a sensitive immunotoxicity indicator. Delayed-type hypersensitivity showed minimal early (6-hour) differences but clear suppression at 24 hours, reflecting impaired T-cell-mediated effector phase inflammation (10).

### Mechanistic Basis: Cellular Profiling via TBNK Analysis

Immunophenotyping via flow cytometry revealed the cellular basis for functional deficits, a novel integration revealing why adaptive responses failed. Cyclophosphamide depleted total lymphocytes, with pronounced B-cell reductions and decreases in CD4⁺ and CD8⁺ T-cells (11,14). B-cells showed particular sensitivity due to cyclophosphamide’s cytotoxicity against proliferating lymphocytes. NK and NKT cells were relatively spared, indicating selective adaptive immune compartment impact. This cellular profiling mechanistically links structural and functional impairments.

### Synthesis of Integrated Endpoints & Risk Interpretation

Hematological changes, lymphoid alterations, suppressed antibody production, attenuated DTH responses, and lymphocyte subset depletion provide mutually supportive evidence of systemic immunosuppression (7,13,15). Consistency across endpoints reduces interpretive uncertainty and enhances biological plausibility. Unlike studies focusing on single endpoints, this coordinated evaluation spans biological organization levels, offering a coherent framework for immune hazard characterization relevant to regulatory toxicology.

## Acknowledgements

The authors sincerely thank Mr. Sai Prasad, CEO of RCC Laboratories, for providing the facilities and support necessary to conduct this study. We are also grateful to Dr. Vishwas K, Sayeed Azgar, and their team for their assistance with histopathology, for the successful completion of this work. This work was supported by RCC Laboratories India Pvt. Limited as an internal project sponsorship.

## Use of AI Assistance

The authors acknowledge the use of AI-based language tools to assist in grammatical corrections and improve clarity of the manuscript. All scientific content, interpretations, and conclusions were independently prepared and verified by the authors.

